# Imperfect integration: sensory congruency between multiple sources modulates selective decision-making processes

**DOI:** 10.1101/2021.03.14.435307

**Authors:** Dominik Krzemiński, Jiaxiang Zhang

## Abstract

Decision making on the basis of multiple information sources is common. However, to what extent such decisions differ from those with a single source remains unclear. Here, we combined cognitive modelling and neural-mass modelling to characterise the neurocognitive process underlying decisionmaking with single or double information sources. Ninety-four human participants performed binary decisions to discriminate the coherent motion direction averaged across two independent apertures. Regardless the angular distance of the apertures, separating motion information into two apertures resulted in a reduction in accuracy. Our modelling results further showed that the addition of the second information source led to a lower signal-to-noise ratio of evidence accumulation with two congruent information sources, and a change in the decision strategy of speed-accuracy trade-off with two incongruent sources. Thus, our findings support a robust behavioural change in relation to multiple information sources, which have congruency-dependent impacts on selective decision-making subcomponents.

## 1 Introduction

Making rapid decisions on the basis of noisy information is a hallmark of voluntary behaviour. Converging evidence from humans (Heekeren et al., 2004) and non-human primates (Roitman and Shadlen, 2002) have supported an evidence accumulation framework governing the decision-making process: the information is integrated over time until the accumulated information in favour of one option reaches a response threshold. This integration process reduces the noise in the accumulated information and facilitates optimal behaviour in terms of accuracy and speed (Bogacz, 2007). A large family of sequential sampling models (Bogacz et al., 2006) can describe adequately the cognitive (Ratcliff and McKoon, 2008) and neural (Wang, 2002) processes during decision making.

Much of research to date on simple decisions has focused on evidence accumulation from a single information source (Gold and Shadlen, 2007). Understanding how a decision-maker integrates information from multiple sources is equally important. In preferential decisions with multiple attributes, such as buying a car based on its colour and price, sequential sampling models can effectively account for various biases and heuristics (Busemeyer et al., 2019), supporting evidence accumulation as a parsimonious decision-making framework for distributed information sources.

The current study focused on another common scenario: making decisions by integrating the *same* type of information originated from multiple sources. For example, when approaching a T junction, a car driver has to consider incoming traffic from both left and right sides of the main road; while entering a roundabout, the driver only need to attend to one side because all vehicles circulate in one direction. An intriguing issue is: how does the presence of multiple information sources affect behavioural performance.

Research on visual search provide circumstantial evidence to imply an imperfect integration of multiple sources, because of the limited capacity of the attentional system. When locating a target among similar distractors or filtering out task irrelevant information, there is a robust behavioural (Palmer, 1995) and neural (Reynolds and Chelazzi, 2004) cost in relation to selective attention on multiple sources. However, two important questions are yet to be addressed. First, when the total amount of information remains unchanged, does separating information into multiple congruent and incongruent sources have the same impact on behaviour? Second, does making decisions with additional information sources lead to a change in the speed of evidence accumulation, the decision threshold, or both?

Here, we addressed these questions in a pre-registered, carefully calibrated online experiment of perceptual decision in two independent groups. Human participants were instructed to decide the average motion direction of random-dot kinematogram from two tilted apertures (Figure 1). Coherent dot motion was presented in both apertures, with their moving directions to be congruent (both leftwards or rightwards) or incongruent (e.g., one leftwards and the other rightwards). In corresponding baseline conditions, coherent motion was presented in a single aperture, with the other aperture containing no coherent motion. Between the two groups, we varied the angular separation of the two apertures, allowing us to evaluate the repeatability of all within-subject effects and assess the between-group effect of aperture angles on behaviour. We fitted a cognitive model, the drift-diffusion model (DDM) (Ratcliff and McKoon, 2008), to behavioural data and inferred the effects of motion congruency and sensory sources on model parameters. Furthermore, we extended a neural-mass model (Wong and Wang, 2006) to demonstrate how the observed behavioural changes can be implemented by a biologically realistic neural network. Together, our study illustrated the neurocognitive mechanisms of perceptual decisions from from multiple sources.

**Figure 1:**
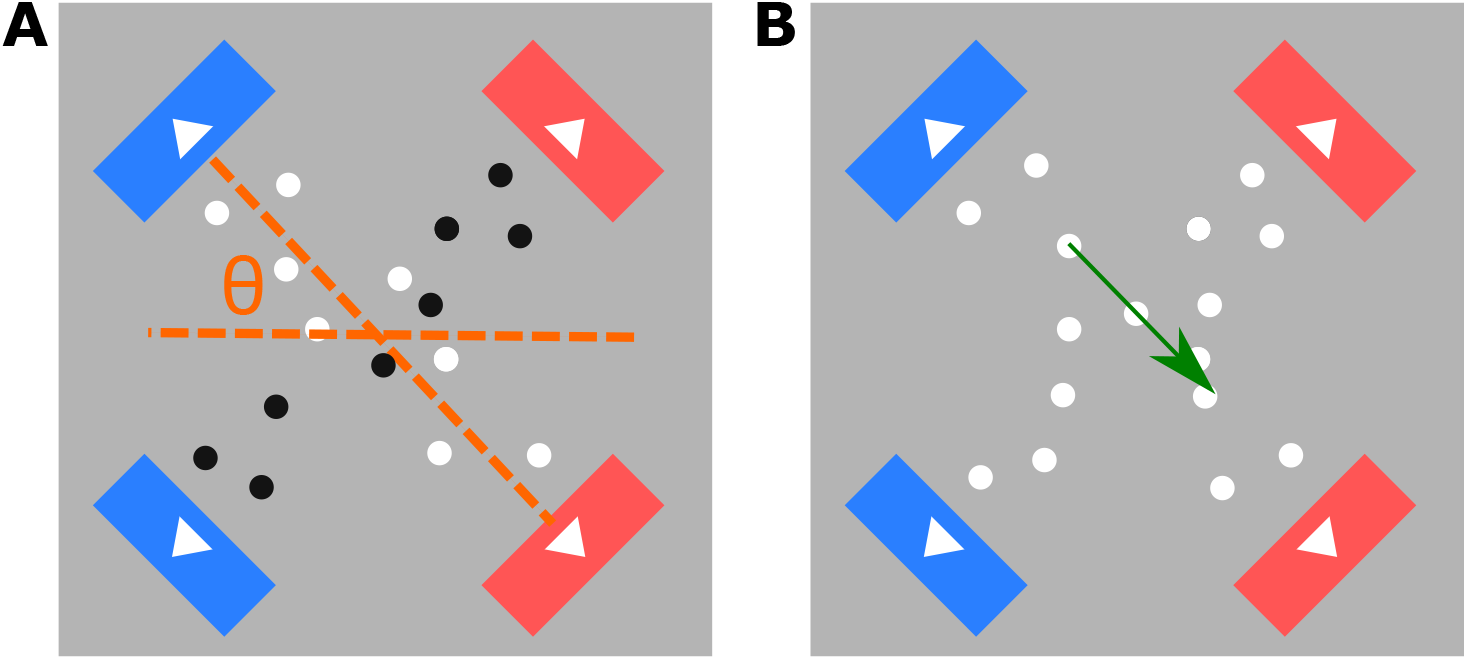
The diagram of the RDK in two rectangular apertures used in (A) the staircase procedure and (B) the main experiment. *θ* denotes the angle between the aperture and the horizontal plane, which was ±20° in Group 1 and ±45° in Group 2. During the staircase procedure, one aperture contained block dots with 0% motion coherence. In the main experiment, both apertures contained white dots.

## 2 Methods

### 2.1 Participants and pre-registration

A total of 94 participants were recruited from an online recruitment portal (*Prolific*, prolific.co) and took part in the experiment online (age range 18-68 years old, median age 25 years old, 25 females, 85 right-handed). Table 1 summaries demographic features of the participants. All participants received monetary payments for their participation. Consent was obtained from all participants. We considered the recruitment from an online portal as a sample of convenience. The study was approved by the Cardiff University School of Psychology Research Ethics Committee.

**Table 1:** Statistical information about participants. NA - data not available, STD - standard deviation.

| Category | Value |
| --- | --- |
| gender | female(25), male(67), NA(1) |
| handness | right(85), left(6), both(2) |
| age (years) | median: 25, mean: 27.3, STD: 8.3 |
| web browser type | Chrome(69), Internet Explorer(14), Firefox(4), Safari(5), NA(2) |
| nationality | United Kingdom(25), Poland(13), Portugal(11), United States(7), Spain(5), Italy(4), Mexico(3), Czech Republic(3), Denmark(2), Ireland(2), Hungary(2), France(2), Lithuania(1), Germany(1), Belgium(1), Sweden(1), Colombia(1), Estonia(1), Finland(1), Netherlands(1), Chile(1), Canada(1), Australia(1), South Africa(1), China(1), NA(1) |

Power analyses, exclusion criteria, experiment procedures and analysis plans were preregistered prior to data collection (https://osf.io/4dn65/). A sample size of *N >* 44 provides > 90% power to detect a medium within-group effect (*d* = 0.5) at *α* = 0.05. We randomly assigned participants into two independent groups. Group 1 (*N* = 49) performed the perceptual decision task with two sources of visual inputs presented along *θ* = ±20°, and Group 2 (*N* = 45) performed the task with visual inputs presented along *θ* = ±45° (see Procedure for details).

### 2.2 Apparatus

The experiment was conducted online. Experimental scripts for stimulus presentation and response collection were written in HTML with a JavaScript library jsPsych 6.0.5 (de Leeuw, 2015) and the jspsych-rdk plugin (Rajananda et al., 2018). The online experiment was hosted on a web server Pavlovia (pavlovia.org), and participants performed the experiment in web browsers on their computers. It has been shown that online experiments in modern web-browsers can serve as a suitable tool for measuring behavioural responses and reaction times with sufficient precision (de Leeuw and Motz, 2016; Semmelmann and Weigelt, 2017; Anwyl-Irvine et al., 2020).

### 2.3 Stimuli

The visual stimuli contained two independently sets of random-dot kinematograms (Britten et al., 1992; Shadlen et al., 1996; Mazurek et al., 2003) displayed within two invisible rectangular apertures (140 pixels width, 550 pixels length) on a grey background (RGB=128, 128, 128). Both rectangular apertures located at the centre of the screen, with one tilted +*θ* from the horizontal plane and the other tilted –*θ*. Hence, the two apertures formed an ‘×’ shape, with *θ* = 20° in Group 1 and *θ* = 45° in Group 2. To facilitate the integration of leftwards and rightward motion across apertures, four motion target indicators were presented at the end of the short edges of the two apertures. On each side of the screen (left or right), the two target indicators had the same colour (red or blue), and the colour assignment of those motion indicators was randomised across participants.

Each rectangular aperture contained 100 dots (i.e., 200 dots in total). Each dot had a radius of 3 pixels. We introduced coherent motion information along the long edge of each aperture (leftwards or rightwards). In each frame, a proportion of dots (namely the motion coherence) was replotted at an appropriate spatial displacement in the direction of motion (2 pixels/frame velocity), relative to their positions in the last frame, and the rest of the dots were replotted at random locations within the aperture. To minimise the impact of local motion information from individual dots, all dots were replotted at random locations after every 7 frames (Rajananda et al., 2018).

### 2.4 Task and procedure

After informed consent and task instructions, the experiment included two parts: (1) a staircase procedure to identify two perceptual thresholds, and (2) the main perceptual decision-making task. In both parts, participants performed a two-alternative forced-choice (2AFC) task, deciding whether the coherent motion direction of the random-dot stimulus is leftward or rightward, either from a single information source (i.e., one aperture in Parts 1, Figure 1A) or combined from double information sources (i.e., two apertures in Part 2, Figure 1B). Participants responded by pressing the ‘k’ key (for leftward decisions) or the ‘p’ key (for rightward decisions) on a keyboard with their right index and middle fingers. Participants took self-paced breaks after each part.

#### 2.4.1 Part 1: staircase procedure

To allow participants to familiarise with stimuli and the task, participants underwent a short practise. The practise part consisted a single block of 32 trials. On each trial, one aperture contained black dots (RGB = 255, 255, 255) with 0% motion coherence, and the other aperture contained white dots (RGB = 0, 0, 0) at one of the four coherence levels (5%, 10%, 20%, and 40%, 8 trials of each level). Participants were instructed to pay attention only to white dots (i.e., the informative aperture) and decide the direction of coherent motion. The coherent motion direction, the order of coherence levels and the informative aperture (i.e., the one at +*θ*° or the one at –*θ*°) were randomised across trials. On each trial, the random-dot stimulus disappeared as soon as a response was made, or a maximum duration of 3500 ms was reached. The inter-trial interval was randomised between 400 and 600 ms.

For online experiments, participants’ hardware settings and their perceptual performance could vary substantially. Therefore, we measured motion discrimination thresholds using the same visual stimulus and the 2AFC task structure as in the practise: one aperture contained black moving dots with 0% (i.e., uninformative) coherence, and the other contained white dots with motion coherence set according to the staircase routine. The direction of coherent motion was randomised across trials. At the end of each trial, visual feedback in text was presented for 500 ms to indicate whether participant’s response was correct or incorrect.

The staircase routine combined two parallel staircase procedures with fixed step sizes: one used a two-down/one-up rule and the other used a three-down/one-up rule. The two staircase procedures are independent and interleaved with each other. In both staircase procedures, the initial motion coherence was set to a supra-threshold value of 31.6%, the ‘up’ step size was 0.1 (log unit) and the ‘down’ step size was 0.074. Therefore, the two-down/one-up procedure converges to the coherence threshold of ~71% accuracy (hereafter referred to as the low coherence *c*_low_, and the three-down/one-up procedure converges to a coherence threshold of ~83% accuracy (hereafter referred to as the high coherence *c*_high_) (Levitt, 1971). Each of the two staircase procedures terminated after ten staircase reversals and the corresponding threshold was calculated as the mean of the motion coherence levels for the last nine reversals.

#### 2.4.2 Part 2: perceptual decisions from from double sources

Part 2 is the main experiment, in which both apertures contained white dots (Figure 1B). Participants were instructed to attend to both apertures and decide whether the coherent motion direction of all (white) dots was leftwards or rightwards.

After task instruction and a brief practise, the main experiment comprised 432 trials, which were divided into 6 blocks of 72 trials. Participants took self-paced breaks between blocks. Decision accuracy (proportion of correct responses) was measured after every two consecutive blocks. If a participant had the accuracy lower than 60%, the experiment ended prematurely and the dataset was discarded from further analysis.

Each block contained 50% of leftwards motion trials and 50% of rightwards motion trials. In each block, 64 main task trials and 8 control trials were presented in a randomised order. The task trials followed a 2 by 2 factorial design with two levels of motion coherence (high and low) and two levels of information sources (single source and double sources).

In the high coherence conditions, trials with double information sources had the low motion coherence *c*_low_ in one aperture and *c*_high_ – *c*_low_ in the other. The coherent motion directions were *congruent* in the two apertures (i.e., both leftwards or both rightwards). Trials with single information source had the high motion coherence *c*_high_ in one aperture and 0% in the other.

In the low coherence conditions, trials with double information sources had the high motion coherence *c*_high_ in one aperture and *c*_high_ – *c*_low_ in the other. Importantly, the coherent motion directions were *incongruent* (i.e., opposite) in the two apertures. Trials with single information source had the low motion coherence *c*_low_ in one aperture and 0% in the other. Therefore, for both double and single information sources, the net motion coherence was always *c*_high_ in high coherence conditions and *c*_low_ in low coherence conditions. In control trials, the motion coherence levels in two apertures was set to 60% and 0%. These easy control trials were served as attention check and excluded from subsequent data analyses.

Each trial started with a 250 ms fixation period, during which a black cross presented in the central of the screen. RDK stimuli in two apertures were then presented for a maximum period of 4000 ms, and the stimuli disappeared as soon as a choice was made. The visual stimulus was followed by an inter-trial interval randomised between 400 and 600 ms.

### 2.5 Data analysis

For the staircase procedure, the non-parametric Mann-Whitney U-test was used to compare the high and low coherence levels (*c*_high_ and *c*_low_) between the two aperture angles (*θ* = 45° and *θ* = 20°). 95% confidence intervals (CI) were obtained using bootstrap procedure with 1000 resamples of simulated distributions.

For the main experiment, we quantified response time (RT) of each trial as the latency between the RDK stimulus onset and behavioural response. To eliminate fast guesses, trials with RT faster than 250 ms were removed. Trials without a valid response were also removed. The discarded trials accounted for 0.26% of all trials. We used mixed frequentist and Bayesian ANOVAs to make group inferences on mean decision accuracy and RT, with the coherence level and the number of information source as within-subject factors. Assumptions of variance equality were checked with Levene’s test. We performed post-hoc comparisons using JASP (http://jasp-stats.org) and used Bayes Factors (*BF*_incl_, *BF*_10_) to characterise the strength of evidence (Wagenmakers et al., 2018).

### 2.6 Cognitive modelling of behavioural data

We used the hierarchical Drift Diffusion Model (DDM) toolbox (Wiecki et al., 2013) to fit DDMs to individual participant’s response time distribution and decision accuracy. The hierarchical DDM assumes that the model parameters of individual participants are sampled from group-level distributions, and the Bayesian fitting procedure estimates the posterior distributions of all model parameters at both individual and group levels, given the observed data.

The basic form of the DDM contained three core parameters (Ratcliff and McKoon, 2008): (1) the drift-rate (*υ*), (2) the decision threshold *a*, and (3) the non-decision time (*T*_er_). For each trial, the model assumes that noisy information is accumulated over time at an averaged rate of *υ* and a starting point of *a*/2, until the accumulated information reaches the upper or the lower decision boundary (*a* or 0) that indicates a correct or incorrect binary response, respectively. The model prediction of RT is the sum of the duration of the accumulation process and the non-decision time, with the latter accounting for delays in sensory encoding and motor execution (Karahan et al., 2019).

To accommodate changes in behavioural performance between conditions, we estimated four variants of the DDM with different parameter constraints. The first three variants allow two of the three parameters (*υ, a, T*_er_) to vary between conditions, and the last variant allows all three parameters to vary. All parameters are allowed to vary between participants in all variants.

For each variant, we generated 20,000 samples from the joint posterior distribution of all model parameters by using Markov chain Monte Carlo sampling. The initial 4,000 samples were discarded for the sake of obtaining stable posterior estimates (Wiecki et al., 2013). To improve the model’s robustness to outliers, we estimated mixture models, in that 95% of the data are explained by the DDM, and 5% of the data are expected to be outliers generated from a uniform distribution (Ratcliff and Tuerlinckx, 2002).

Model fits were assessed by comparing each model’s deviance information criterion (DIC) value (Spiegelhalter et al., 2002), which takes into account both the log-likelihood function of observed data and the complexity of the model. For the best fitting variant, we used Bayesian hypothesis testing (Gelman et al., 2013) to make inferences between conditions from the parameters’ group-level posterior distributions. For consistency, we use *p* to refer to frequentist *p*-values, and *P_p_*|*_D_* to refer to the proportion of posteriors supporting the testing hypothesis at the group level from Bayesian hypothesis testing.

### 2.7 Recurrent neural mass model

We further used a neural mass model (Wong and Wang, 2006) to qualitatively demonstrate the effects of motion coherence and the number of information source on behavioural performance. The model considered here is simplified from a recurrent spiking neural network model (Wang, 2002) via the mean-field approximation. Specifically, the neural-mass model includes two neural populations (i.e., accumulators) supporting two alternatives in a decision-making task. The neural accumulators have self-excitatory and mutual inhibitory connections. Each accumulator receives selective external inputs (*I*_in,L_ and *I*_in,R_) as momentary evidence supporting each alternative (e.g. leftwards vs rightwards motion), as well as a common, non-selective background input *I*_0_ (Figure 7A). During decision making, two accumulators compete against each other, and the first accumulator that reaches a decision threshold renders the corresponding response. It has been shown that this biologically realistic neural mass model can explain behavioural and single-unit recording data from perceptual decision experiments (Wong and Wang, 2006). Moreover, within a certain parameter range, the dynamics of the model can mathematically approximate that of the DDM (Wong and Wang, 2006; Bogacz et al., 2006).

**Figure 7:**
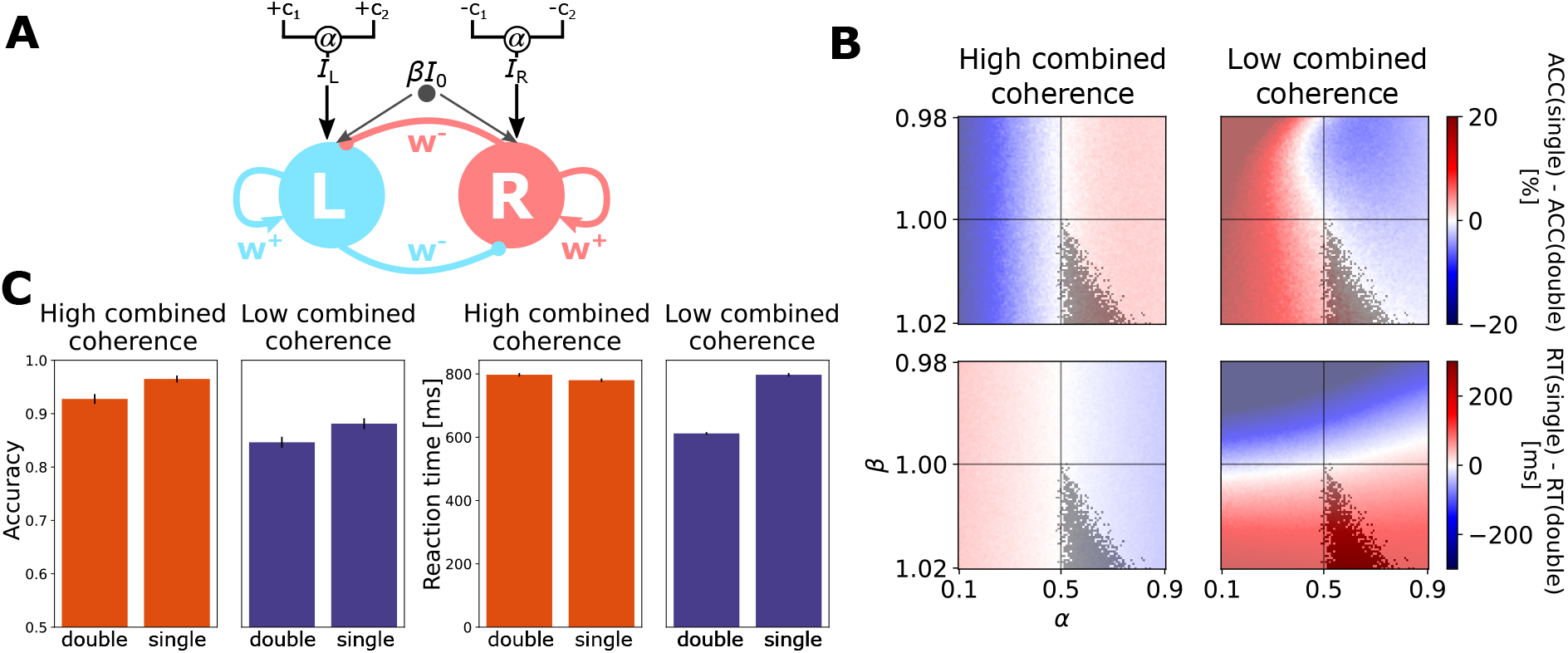
Neural-mass model simulation results. (A) The diagram of the two-state neural-mass model. *w*^+^ denotes excitatory connections, *w*^-^ denotes inhibitory connections, *α* determines how the weight of sensory evidence is split between two sources, and *β* modulates non-selective background input current. (B) Parameters space with difference in performance between single and double informatino sources in high combined evidence (left) and low combined evidence (right) conditions in terms of accuracy (top row) and reaction times (bottom row). Grey zone indicates area where the model parameters reproduce the direction of behavioural differences observed in the experiment. *α* varied between 0.1 and 0.9, and *β* varied between 0.98 and 1.02. (C) Behavioural performance from model simulations with parameters *α* = 0.7 and *β* = 1.018.

Here, we extended the original neural-mass model to take into account the presence of the two information sources in the current study (for modelling details see Supplementary methods). The deterministic input currents (*I*_in,L_ and *I*_in,R_) to the two neural accumulators are given by

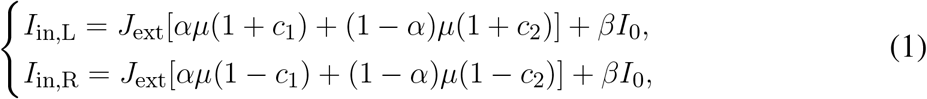

where the first term is the selective input current and the second represents the non-selective background input current. Here, *c*_1_ and *c*_2_ denote the motion coherence levels in the two independent apertures. For simplicity, hereafter we assign *c*_1_ to represent the stronger coherence between the two (|*c*_1_| > |*c*_2_|). Other parameters were set in line with previous studies (Wong and Wang, 2006; Standage et al., 2014b): *J*_ext_ = 5.2 · 10^−4^ nA · Hz^−1^ is the the average synaptic coupling parameter, *I*_0_ = 0.32lnA represents the baseline of the background input current, and *μ* = 35 Hz is the baseline of input strength of the evidence.

The two scaling parameters *α* and *β* control to what extent task conditions affect model inputs. First, in trials with double informative sources, participants need to combine the evidence from two apertures for optimal decisions. The parameter *α* determines how a decision maker splits the weight of sensory evidence from two sources. *α* = 0.5 implies that a participant weight two sources equally, while 0.5 < *α* < 1 or 0 < *α* < 0.5 implies that the dominant source is weighted more or less, respectively.

Second, previous studies suggest that a change in the baseline input *I*_0_ results in speed-accuracy trade-off (Standage et al., 2014b; Heitz and Schall, 2012). Compared with the single source condition, the double source condition with incongruent motion directions had lower accuracy and faster RT, suggesting that participants may trade accuracy for speed in the presence of conflict information. Therefore, we assumed that the non-selective background input current is modulated by a factor of *β* in that condition, which changes the model dynamics and in turn affects both the accuracy and RT relative to the condition with a single information source. For other conditions, we set *β* =1 such that the non-selective input is at its baseline level.

To identify the parameter regime where the neural mass model can produce qualitatively the behavioural pattern observed in the experiment, we ran model simulations with different values of *α* and *β*. For each parameter set, we ran 5,000 simulations of each of the four experimental conditions with representative coherence levels (*c*_high_ = 20% and *c*_low_ = 15%). For example, in the incongruent condition with double information sources, *c*_1_ = 20% and *c*_2_ = –5% for leftward motion; and *c*_1_ = –20% and *c*_2_ = 5% for rightward motion. Mean accuracy and RT of each condition was then calculated from all simulations.

### 2.8 Open data and scripts

We have made the data (https://figshare.com/articles/dataset/13567916), all analyses scripts and experimental materials (https://github.com/dokato/2drdk) open access.

### 3 Results

#### 3.1 Behavioural results

Two groups of participants performed a coherent motion discrimination task, with independent RDK stimuli in two apertures at an angle of 20° (Group 1) or 45° (Group 2) (Figure 1A). Prior to the main experiment, all participants underwent a fixed-size staircase procedure to estimate two motion coherence thresholds: *c*_low_ from a two-down/one-up rule and *c*_high_ from a three-down/one-up rule (Figure 2 and Supplementary Figure 1). A Mann Whitney *U* test showed no significant difference in coherence thresholds between the two groups with different aperture angles (*c*_low_: *U*(45, 49) = 983, *p* = 0.18, 95%CI = [–2.7,0.9]%; *c*_high_: *U*(45, 49) = 952, *p* = 0.13, 95%CI = [–5.4, 1.3]%). As expected, the coherence threshold *c*_high_ was significantly larger than *c*_low_ (*U*(94) = 1534, *p* < 0.0001, 95%CI = [–9.1, –5.1]%). These results suggest that participants achieved reliable performance in the motion discrimination task online.

**Figure 2:**
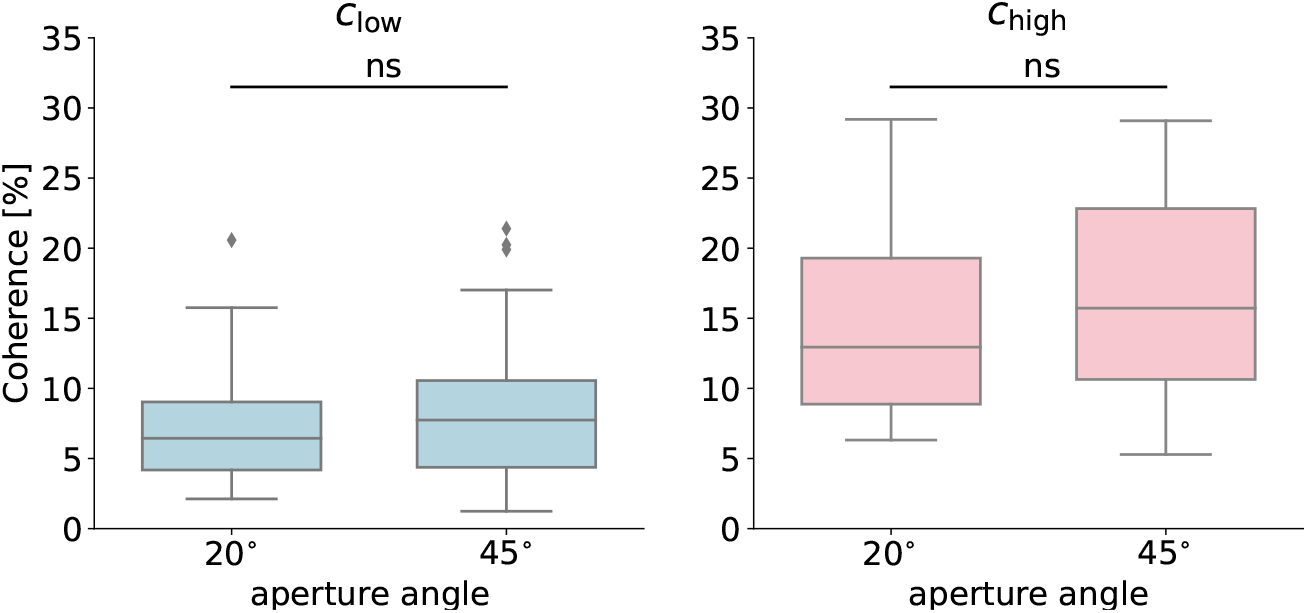
Staircase procedure results. Two motion coherence thresholds obtained from a parallel staircase routine: *c*_low_ from the two-down/one-up rule and *c*_high_ from the three-down/one-up rule. There were no significant (ns) differences between coherence values for the aperture angles *θ* = 20° and *θ* = 45°.

In the main experiment, participants in both groups decided the combined coherent motion direction (leftwards vs. rightwards) in a 2-by-2 factorial design: either single or double apertures contained non-zero motion coherence, and the combined coherence level in the two apertures was either *c*_low_ or *c*_high_ (Figure 1B). We quantified participant’s performance in mean decision accuracy (proportion of correct) and RT.

The two groups with different angles of stimulus apertures achieved similar performance. A two-way mixed ANOVA showed no significant group effect on behavioural performance (accuracy: *F*(1, 92) = 0.009, *p* = 0.92, 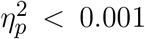, BF_incl_ = 0.29; RT: *F*(1, 92) = 0.05, *p* = 0.30, 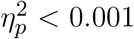, BF_incl_ = 0.24). The participant’s grouping interacted with combined coherence levels for accuracy: (*F*(1, 92) = 4.27, *p* = 0.04, 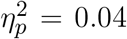, BF_incl_ = 4.20), but not RT: *F*(1, 92) = 1.04, *p* = 0.31, 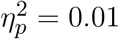, BF_incl_ = 0.19), nor the number of information sources (accuracy: *F*(1, 92) = 0.02, *p* = 0.88, 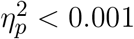, BF_incl_ = 0.17; RT: *F*(1, 92) = 0.01, *p* = 0.93, 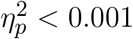, BF_incl_ = 0.15).

Across both groups, the high combined coherence *c*_high_ led to better accuracy (Figure 3, *F*(1, 92) = 269.25, *p* < 0.001, 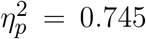, BF_incl_ = 2.3 · 10^55^) and faster RT (Figure 4, *F*(1, 92) = 53.70, *p* < 0.001, 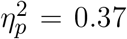, BF_incl_ = 3.73) than the low combined coherence *c*_low_. Compared with conditions of single information source, splitting motion information into two apertures resulted in lower accuracy (*F*(1, 92) = 47.50, *p* < 0.001, 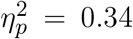, BF_incl_ = 1.7 · 10^4^) without a significant main effect on RT (*F*(1, 92) = 0.14, *p* = 0.71, 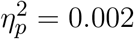, BF_incl_ = 0.12).

**Figure 3:**
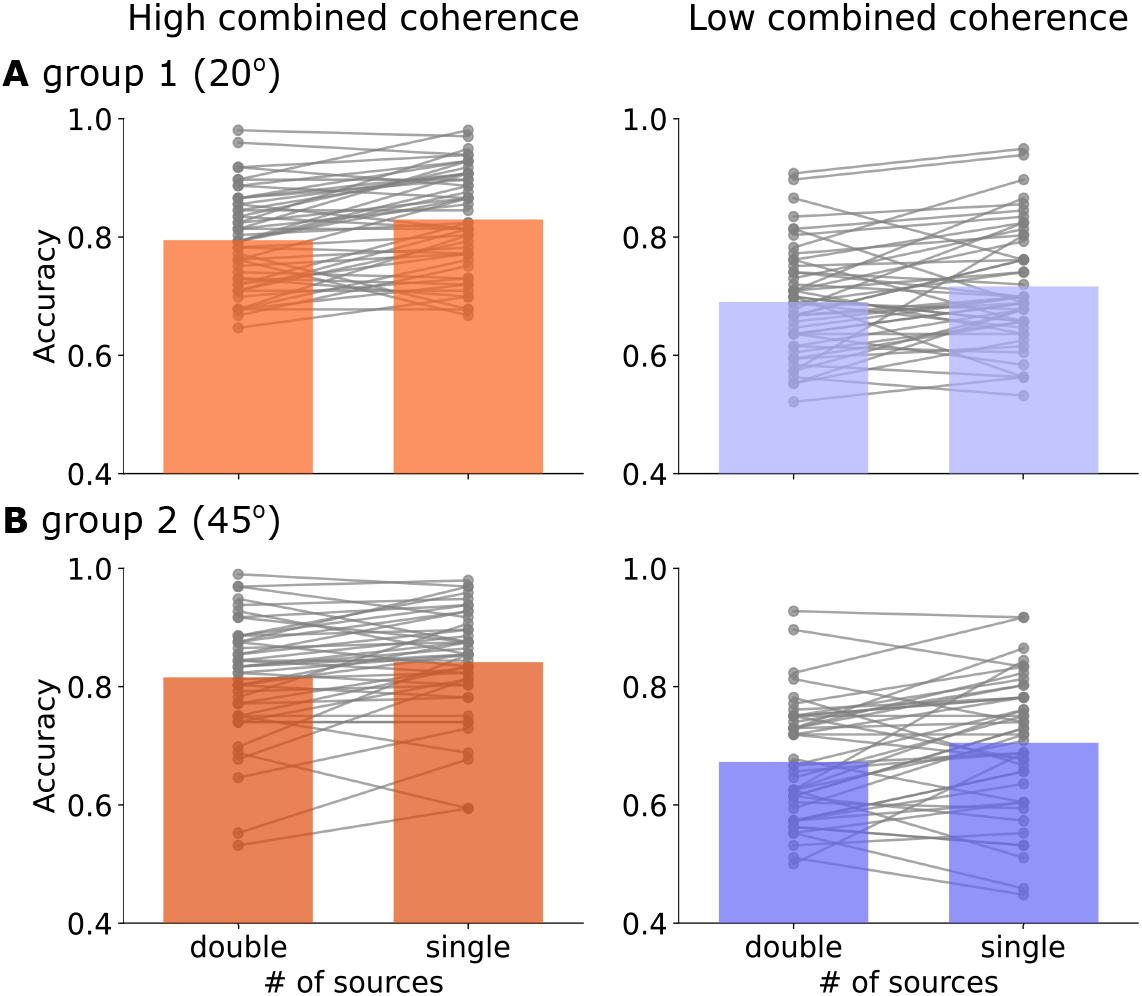
Accuracy (proportion of correct responses) in the main experiment for high (red) and low (purple) combined evidence condition. Bars represent the averaged accuracy in (A) Group 1 (aperture angle *θ* = ±20°) and (B) Group 2 (*θ* = ±45°). Grey dots represent individual participants’ accuracy. Each solid line links the performance between double and single source conditions from the same participant.

**Figure 4:**
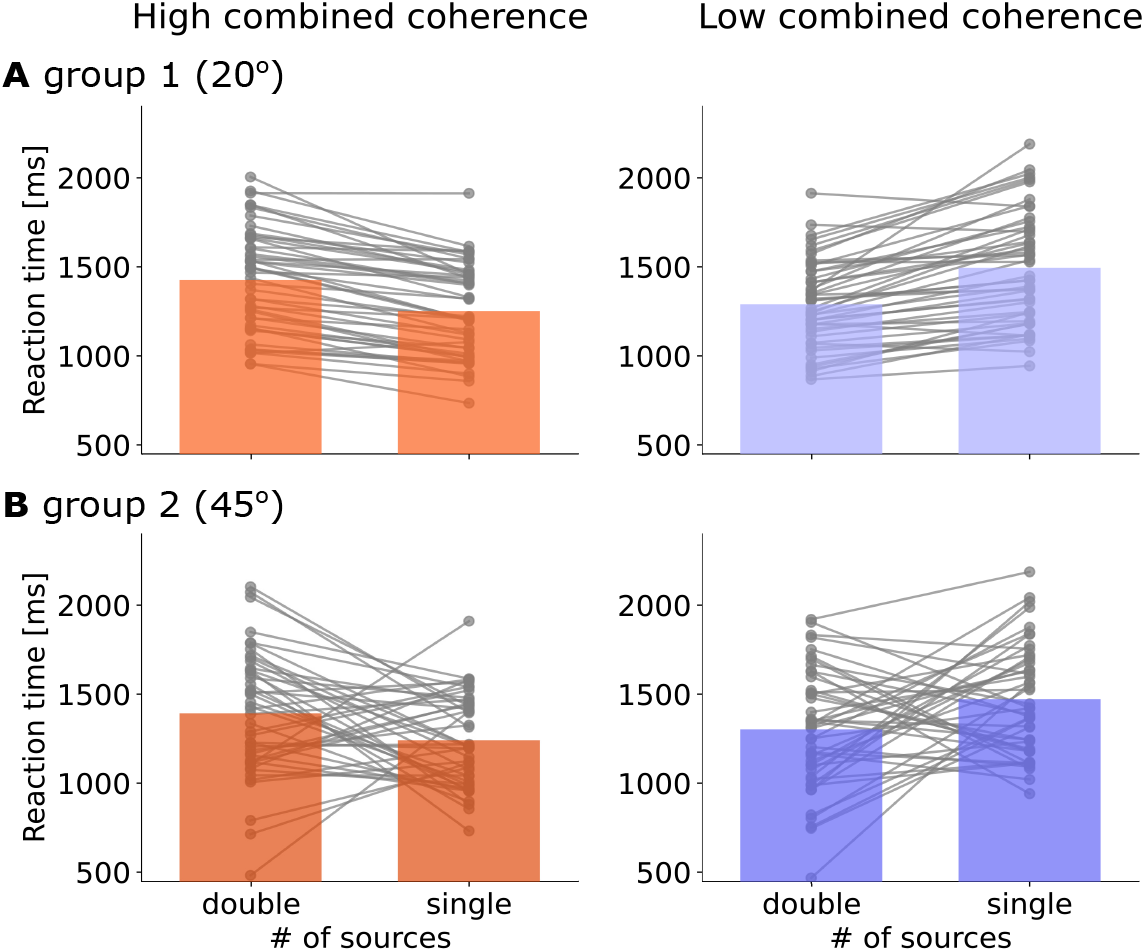
Reaction time (RT) in the main experiment for high (red) and low (purple) combined evidence condition. Bars represent the averaged accuracy in (A) Group 1 (aperture angle *θ* = ±20°) and (B) Group 2 (*θ* = ±45°). Grey dots represent individual participants’ accuracy. Each solid line links the performance between double and single source conditions from the same participant.

There was no interaction in accuracy between combined coherence levels and the number of information sources (*F*(1, 92) = 0.01, *p* = 0.92, 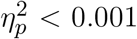, BF_incl_ = 0.17). For RT, the interaction between the two factors was significant (*F*(1, 92) = 208.38, *p* < 0.001, 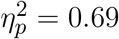, BF_incl_ = 1.2 · 10^13^), suggesting that presenting motion information in double apertures elicited different changes in response speed between the two combined coherence levels (Figure 4). It is worth noting that in *c*_high_ conditions, motion directions in two apertures were congruent in the case of double information sources (i.e., both leftwards or both rightwards). A post-hoc test showed that compared with single source conditions, congruent double source conditions had slower RT (BF_10_ = 8014.01, Bayesian *t*-test). Conversely, in *c*_low_ conditions, motion directions in two apertures were incongruent (e.g., one leftwards and the other rightwards) and led to faster RT (BF_10_ = 1.21 · 10^4^).

#### 3.2 Cognitive modelling results

We used a hierarchical Bayesian implementation (Wiecki et al., 2013; Vandekerckhove et al., 2011) of the DDM (Ratcliff, 2002; Bogacz et al., 2006) to decompose individual participant’s accuracy and RT into model parameters that quantify latent cognitive processes. We considered four model variants, which allow the drift-rate *υ*, the non-decision time *T*_er_ and the decision threshold *a* to be fixed or vary between task conditions.

For each model variant, the Gelman-Rubin *Ȓ* convergence criterion (Gelman and Rubin, 1992) was used to assess the convergence of the last 16,000 MCMC samples from 5 independent Markov chains. The maximum value of the statistic from all parameters was *Ȓ* = 1.0012, which is lower than the criterion of convergence 1.1 (Gelman and Rubin, 1992), suggesting that all parameter estimates converged after 20,000 steps.

The model variant that described the data best (i.e., the one with the lowest DIC value) allows all three parameters (*υ, T*_er_ and *a*) to vary between conditions. To evaluate the model fit, we generated model predictions by simulations with the posterior estimates of the model parameters. There was a good agreement between the observed data and the model simulations in all conditions (Figure 6 and Supplementary Figures 2 and 3).

**Figure 6:**
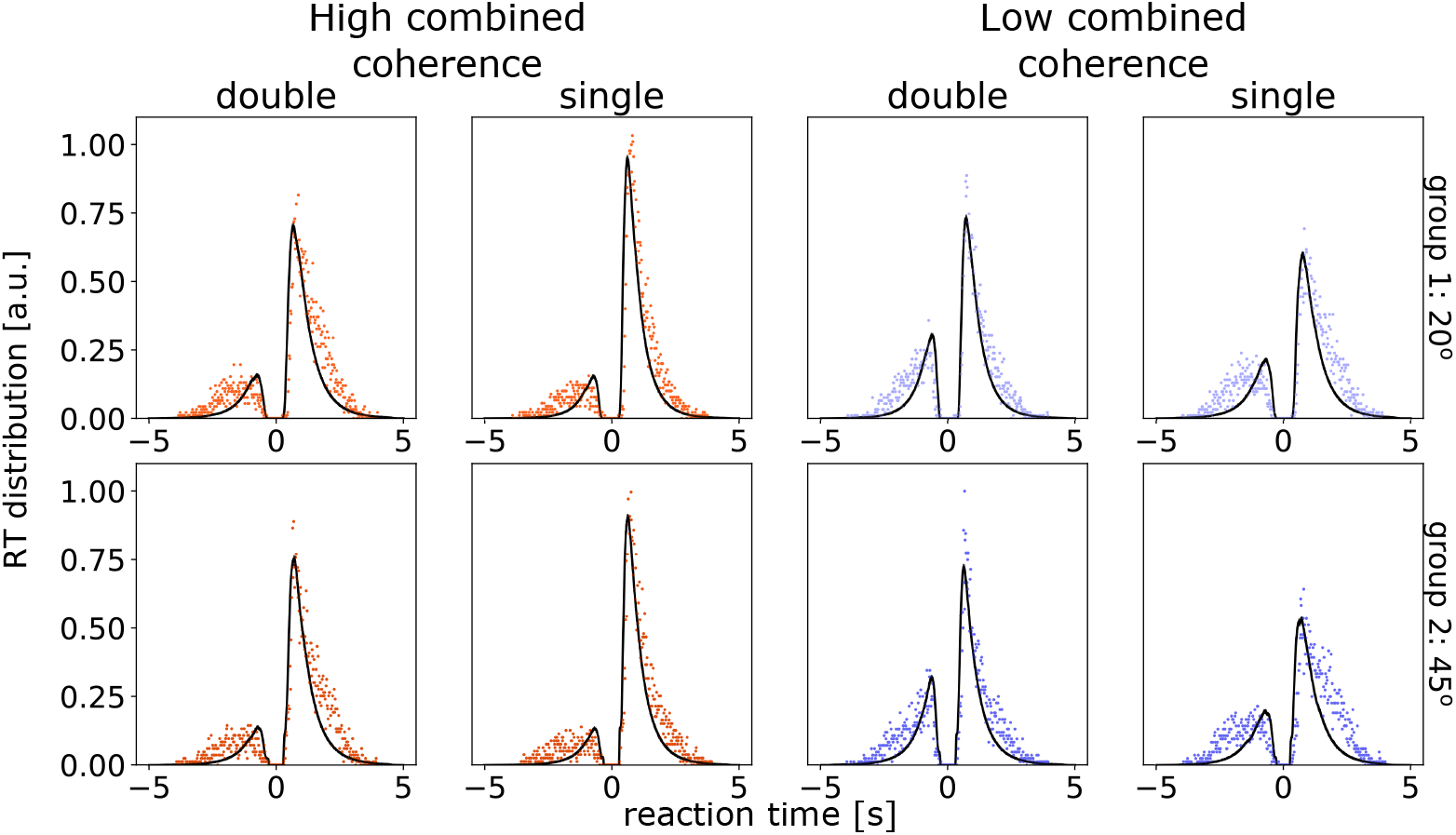
Posterior predictive response time (RT) distributions from the fitted DDM. Each panel shows normalised histogram of the observed data (red for high combined coherence and blue for low combined coherence conditions) and the model predictions (black lines) across participants. The RT distribution of correct responses is shown along the positive horizontal axis. The RT distribution of error responses is shown along the negative horizontal axis. The posterior predictions of the model were generated by averaging 1000 simulations of the same amount of observed data. Top row shows results from Group 1 (with 20° angle between apertures) and the bottom row for Group 2 (45°).

Figure 5C shows the posterior parameter estimates for the two participant groups. We used Bayesian statistics (Gelman et al., 2013; Kruschke, 2014) to quantify the proportion of parameters’ posterior distributions that did not overlap between groups and conditions (Table 2). There was no evidence to support a difference in model parameters between groups (*P_p_*|*_D_* < 0.93 in all parameters). For the drift-rate, there were strong evidence to support differences between all conditions (Group 1: *P_p_*|*_D_* > 0.998; Group 2: *P_p_*|*_D_* > 0.989, Table 2), except between the single and double source (i.e., the incongruent condition) conditions with the low coherence level (Group 1: *P_p_*|*_D_* = 0.662; Group 2: *P_p_*|*_D_* = 0.671). The incongruent condition with double information sources had a lower decision threshold than the other three conditions (Group 1: *P_p_*|*_D_* > 0.958; Group 2: *P_p_*|*_D_* > 0.984). We did not observe strong evidence in supporting a difference in the non-decision time between conditions (Group 1: *P_p_*|*_D_* < 0.71; Group 2: *P_p_*|*_D_* < 0.74).

**Figure 5:**
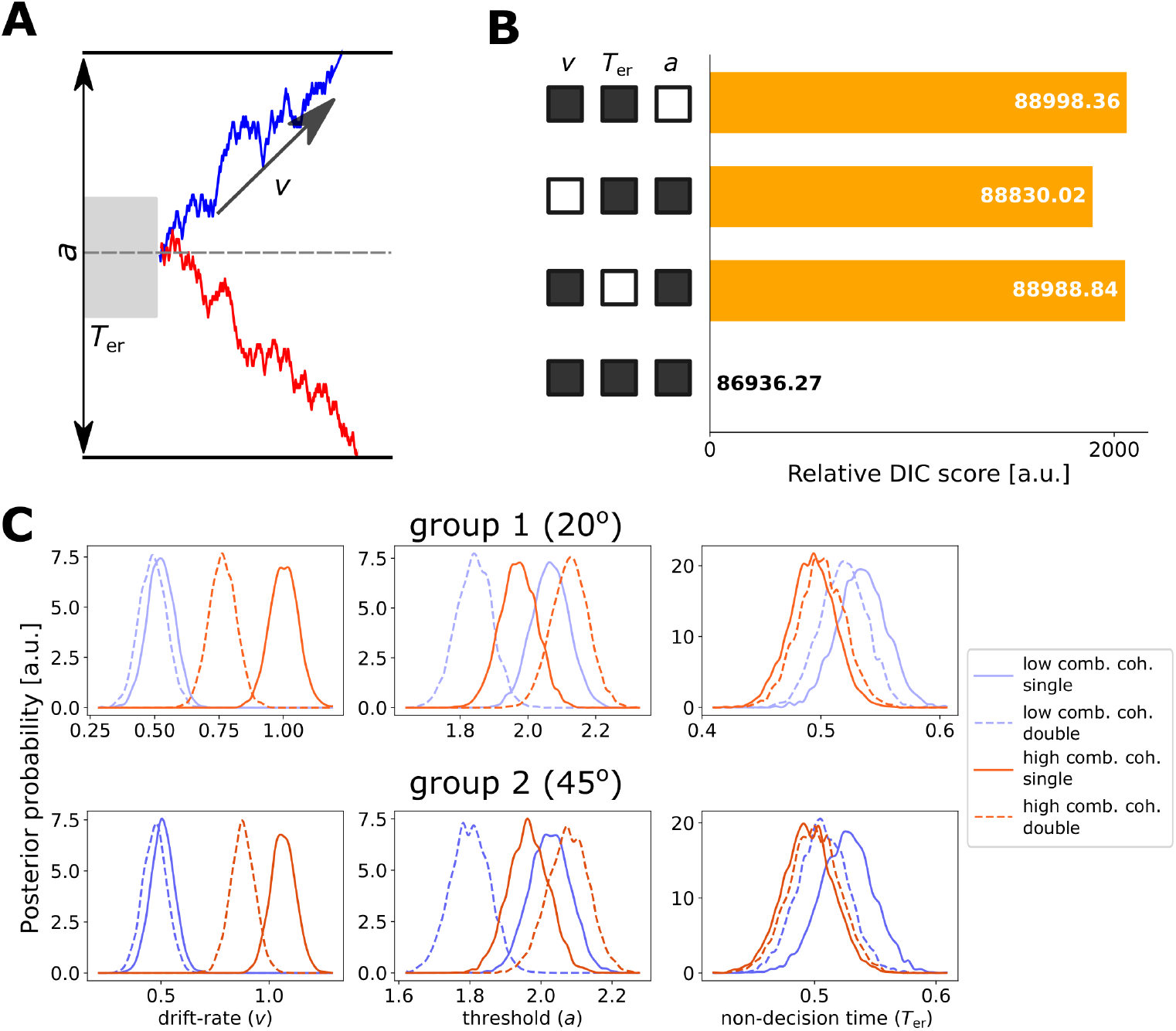
Drift-Diffusion Model (DDM) fitting results. (A) Examples of evidence accumulation trajectories depicted by the DDM. The decision threshold *a* represents the distance between the correct and incorrect decision thresholds. The drift-rate *υ* describes the average speed of evidence accumulation. The non-decision time *T*_er_ represents the latency of other processes not included in the evidence accumulation. The diffusion continues until the accumulated evidence reaches one of the two thresholds (solid black lines). If the accumulated evidence reaches the correct (upper) threshold (blue trajectory), the model predicts a correct response. Because of noise, the accumulated evidence may reach the incorrect (lower) threshold (red trajectory). (B) The deviance information criterion (DIC) value differences between the four variants of the DDM and the best fit. The black square indicates that the corresponding parameter can vary between the conditions, and the white square indicates that the parameter is invariant. The best model had variable *a, υ* and *T*_er_ between conditions. (C) The posterior distributions of parameter values of the best fit model (top: Group 1 with 20° aperture angle); bottom: Group 2 with 45° aperture angle.

**Table 2:** Posterior comparisons of model parameters. The table listed the proportion of non-overlap between two posterior parameter estimates *x* and *y*, which is equivalent to a Bayesian test of the hypothesis *P_p_*|*_D_*(*x* > *y*). Experimental conditions: I1 and I2 refer to single or double informative sources with a combined coherence of *c*_high_; C1 and C2 refer to single or double informative sources with a combined coherence of *c*_low_. The DDM model parameters: *υ* dirft-rate, *a* decision threshold and *T*_er_ non-decision time. Two angular distances in two groups: *θ*_1_ = 20°, *θ* = 45°.

| | $v$ | | $a$ | | $T_{\text{er}}$ | |
| --- | --- | --- | --- | --- | --- | --- |
| | $\theta_1$ | $\theta_2$ | $\theta_1$ | $\theta_2$ | $\theta_1$ | $\theta_2$ |
| x = I1; y = I2 | 0.662 | 0.671 | 0.997 | 0.998 | 0.708 | 0.744 |
| x = C1; y = I2 | 1.000 | 1.000 | 0.958 | 0.984 | 0.173 | 0.339 |
| x = C2; y = I2 | 1.000 | 1.000 | 0.999 | 0.999 | 0.250 | 0.402 |
| x = C1; y = I1 | 1.000 | 1.000 | 0.113 | 0.233 | 0.067 | 0.145 |
| x = C2; y = I1 | 0.999 | 1.000 | 0.770 | 0.734 | 0.118 | 0.186 |
| x = C1; y = C2 | 0.998 | 0.989 | 0.025 | 0.087 | 0.392 | 0.430 |

#### 3.3 Neural-mass modelling results

Our cognitive modelling results suggested that splitting coherent motion information into two apertures led to a decrease of drift-rate in the congruent condition, and a decrease of decision threshold in the incongruent condition. How could these changes be incorporated in a biologically realistic model? To address this question, we introduced two extensions (Figure 7A) to a neural-mass model of perceptual decision (Wong and Wang, 2006), which implements an evidence accumulation process akin to that of the DDM (Bogacz et al., 2006). First, for conditions with double information sources, we assumed that the sensory input selective to motion coherence is a weighted sum of the two sources. The two weights (*αμ* and (1 – *α*)*μ*; see Equation 1) sum up to the constant baseline weight *μ* that is applied to the conditions with single information source. Second, we assumed that the non-selective sensory input *I*_0_ is changed at the rate of *β* in the double source condition with incongruent inputs, which has been shown to be a realistic neural mechanism in modulating decision threshold (Heitz and Schall, 2012; Standage et al., 2014a).

We parametrically modulated the two scaling parameters *α* and *β*. For each parameter set, we simulated the extended neural-mass model 20,000 trials (5,000 simulations for each experimental condition) and estimated the decision accuracy as well as mean RT. Figures 7B and 7C show the behavioural performance from simulations. We further identified parameter regimes that qualitatively satisfy the observed performance difference between double (both congruent and incongruent) and single information sources conditions. Based on model simulations, the scaling parameter *α* on input weights needs to be larger than 0.5, suggesting that the dominant, or more informative, sensory input of the two apertures is weighted more than the other. The parameter *β* needs to be larger than 1, suggesting that incongruent double information sources are associated with an elevated non-selective sensory input.

## 4 Discussion

The current study examined, in two independent groups, how the presence of a second source of sensory information affects the behavioural performance of perceptual decision as well as its underlying neurocognitive mechanisms. When motion directions are congruent between the two sources, decisions on the global motion direction were less accurate and slower than that in the single-source condition with the same amount of total information (i.e., combined motion coherence). In contrast, when two information sources are incongruent, decisions were less accurate but faster than that in the single-source condition. Therefore, the change in task performance depends on the congruency between multiple sources of sensory evidence.

Using a Bayesian DDM, our cognitive modelling provided novel evidence on the decisionmaking process with multiple information sources. First, information congruency has selective influence on different decision-making subcomponents. The congruent, double-source condition had a lower drift-rate than its corresponding single-source condition (i.e., with a combined motion coherence of *c*_high_ in both). The drift-rate of the DDM represents the signal-to-noise ratio of the information (Ratcliff and McKoon, 2008) and has been linked to the allocation of attention (Schmiedek et al., 2007). The presence of congruent information in two apertures may modulate the divided attention towards the stimuli that in turn lowers the averaged rate of evidence accumulation. Human electrophysiological data support this proposition. For two peripheral visual patches presented simultaneously, the early-visual ERP component is characteristic to the attended location, as its amplitude is maximal over posterior electrodes contralateral to the attending side (Eimer, 1996; Luck and Hillyard, 1994). Recent studies showed that this EEG marker of selective attention modulates the rate of evidence accumulation in perceptual decision (Loughnane et al., 2016), and the dynamics of selective attention can influence evidence accumulation throughout the decision process (Rangelov and Mattingley, 2020).

Second, splitting motion information into two incongruent apertures did not vary the drift-rate. Instead, there was a substantial reduction in the decision threshold, reflecting the behavioural change that participants traded accuracy for speed in this condition. The speed-accuracy trade-off (SAT) is widely observed across decision-making tasks (Wickelgren, 1977; Heitz, 2014; Beersma et al., 2003). In experiments with humans, the SAT is often induced explicitly via verbal instructions (Zhang and Bogacz, 2010) or response deadlines (Yamaguchi et al., 2013). Such manipulations can efficiently switch between accuracy-seeking and speed-seeking behaviour every few trials (Mulder et al., 2013) or in consecutive trials (Forstmann et al., 2008). Modelling studies on explicit SAT demands have been consistently associated with the change of decision threshold (Palmer et al., 2005; Ratcliff, 2006): a smaller decision threshold leads to faster and more error-prone decisions. Nevertheless, the SAT can also be triggered endogenously without explicit demands (Desender et al., 2019). In the current study, the two apertures in the incongruent condition contained contradictory information, presenting a decision dilemma. Our results showed that, in such difficult scenario, participants adapted their decision strategy to be more speed-seeking, allowing them to complete the current decision sooner. Future research could examine this conflict avoidance bias further, by changing the relative difference between multiple incongruent information sources.

Third, it is worth comparing between single- and double-source conditions which had equal motion coherence in the dominant aperture. Compared with the single-source condition with high coherence (*c*_high_ in one aperture and 0% in the other), the incongruent double-source condition (*c*_high_ in one aperture and *c*_high_ – *c*_low_ in the other) condition had a smaller drift-rate. The congruent double-source condition had a larger drift-rate than the single-source condition with low coherence. That is, introducing additional incongruent (or congruent) information led to a reduction (or increase) in the rate of evidence accumulation. These results agree with two robust behavioural effects consistently reported in the literature of visual search: the presence of distractors in hindering the search performance (Palmer, 1995), as well as the facilitating role of task relevant information (Krummenacher et al., 2002). Our findings further support that participants did not only attend the dominant aperture, but attempt to integrate motion information across apertures to form decisions, albeit the integration of multiple information sources was not optimal, as discussed above and reported elsewhere (Wyart et al., 2015).

Fourth, the *T*_er_ is considered as the latency external to the evidence accumulation process (Ratcliff and McKoon, 2008). Recent electrophysiological and imaging studies suggest that the *T*_er_ accounts for delays in early sensory processing (Nunez et al., 2019) or motor preparation (Karahan et al., 2019). The current study did not observe a change in the *T*_er_ between task conditions in either participant group. Hence, our results are unlikely originated from potential changes in early visual processing or motor execution in response to multiple information sources.

Based on our cognitive modelling, we proposed two extensions to a neural-mass model of decision-making (Wong and Wang, 2006). The first extension is to vary the baseline weights between sensory inputs from two independent apertures, and the second is to vary the non-selective background inputs in the incongruent double source condition. From an exhaustive search of the parameter space, we identified the parameter regime that can qualitatively account for the observed behavioural changes in the presence of two information sources. It is worth noting that the neural-mass model is not meant to fit to experimental data, but provides a biologically plausible interpretation of their neural implementations.

We showed that, to accommodate experimental results, the sensory input from the dominant source needs to be weighted higher than the input from the additional source (*α* > 0.5). When this ratio becomes too high, the contribution of the additional source diminishes, resulting in the model unable to integrate information from the non-dominant source. Therefore, perceptual decisions with two information sources involve an unbalanced integration that is biased towards the more informative source.

Additionally, the non-selective background input needs to be elevated in the incongruent double-source condition (*β* > 1). An increased baseline activity effectively decreases the amount of evidence required to reach a decision threshold (Standage et al., 2014a), leading to speed-seeking behaviour at the cost of less accurate decisions that was observed in the current study. Both brain imaging (Ivanoff et al., 2008) and single-unit recording (Heitz and Schall, 2012) studies showed that the baseline change underlies the SAT, consistent with our model simulation results.

Interestingly, although participants were instructed to decide leftwards vs. rightwards coherent motion from two tilted apertures, the angular distance between the apertures did not affect behaviour nor DDM parameters. This may seem counterintuitive, because a larger angular distance results in less coherent motion information to be projected onto the horizontal plane. Future studies could examine whether there is a significant behavioural difference at larger aperture angles, because, in an extreme condition of two vertical apertures (*θ* = ±90°), there is zero horizontal motion and the decision accuracy will be at chance. One plausible account for the lack of group difference is that participants decided the coherent motion direction with a reference of individual apertures (i.e., along their long edges), not the horizontal plane. One could validate this hypothesis by presenting multiple independent sources of motion information within a single aperture (e.g. (Wendelken et al., 2009)).

There are several limitations of this study. First, as in all online experiments, the current study faced practical constraints that could affect the millisecond-level precision of stimulus timing (Anwyl-Irvine et al., 2020). To mitigate the impact of variable testing environments between participants, we pre-registered the experiment, applied rigorous inclusion/exclusion criteria, conducted staircase procedures to calibrate stimuli for individual participants, and focused on within-subject effects in most analyses. Our study and research practises contribute to the growing trend of online psychological, or even psychophysical experiments, confirming the feasibility and reproducibility (i.e., in two independent groups) of online experiments to investigate task-specific effects in the context of perceptual decision-making (Semmelmann and Weigelt, 2017; de Leeuw and Motz, 2016).

Second, owing to the potential variability of online testing environments between participants, we designed our experiment to be completed in one testing session. Perceptual learning studies showed that behavioural performance of coherent motion discrimination improves steadily over multiple testing sessions across several days (Zhang and Rowe, 2014; Liu and Watanabe, 2012). It would be of interest to examine if repetitive training modulates the behavioural change between single and multiple information sources.

In conclusion, when sensory information is separated into independent apertures, perceptual decisions are less accurate. Our cognitive and neural-mass modelling showed two selective neurocognitive mechanisms underlying the behavioural effect, a change in the signal-to-noise ratio of the accumulation process and the speed-accuracy trade-off, depending on the congruency of multiple sensory sources. These findings suggest that both attentional demands and endogenous response strategies influence flexible decision-making in humans.

## Author contributions

DK: conception and design, data analysis, computer simulations, interpretation of results, drafting and revising the manuscript; JZ: conception and design, supervision of data analysis, interpretation of results, revising the manuscript.

## Acknowledgements

DK was supported by a Engineering and Physical Sciences Research Council PhD Scholarship (EP/N509449/1). JZ was supported by European Research Council (716321).

## References

Anwyl-Irvine A, Dalmaijer ES, Hodges N, and Evershed JK (2020). Realistic precision and accuracy of online experiment platforms, web browsers, and devices. Behavior Research Methods, pages 1—19.

Beersma B, Hollenbeck JR, Humphrey SE, Moon H, Conlon DE, and Ilgen DR (2003). Cooperation, competition, and team performance: Toward a contingency approach. Academy of Management Journal, 46(5):572—590.

Bogacz R (2007). Optimal decision-making theories: linking neurobiology with behaviour. Trends in cognitive sciences, 11(3):118—125.

Bogacz R, Brown E, Moehlis J, Holmes P, and Cohen JD (2006). The physics of optimal decision making: A formal analysis of models of performance in two-alternative forced-choice tasks. Psychological Review, 113:700—765.

Britten K, Shadlen M, Newsome W, and Movshon J (1992). The analysis of visual motion: A comparison of neuronal and psychophysical performance. Journal of Neuroscience, 12(12):4745–4765.

Busemeyer JR, Gluth S, Rieskamp J, and Turner BM (2019). Cognitive and neural bases of multi-attribute, multi-alternative, value-based decisions. Trends in Cognitive Sciences, 23(3):251–263.

de Leeuw JR (2015). jsPsych: A javascript library for creating behavioral experiments in a web browser. Behavior Research Methods, 47(1):1–12.

de Leeuw JR and Motz BA (2016). Psychophysics in a web browser? Comparing response times collected with javascript and psychophysics toolbox in a visual search task. Behavior Research Methods, 48(1):1–12.

Desender K, Boldt A, Verguts T, and Donner TH (2019). Confidence predicts speed-accuracy tradeoff for subsequent decisions. Elife, 8:e43499.

Eimer M (1996). The n2pc component as an indicator of attentional selectivity. Electroencephalography and clinical neurophysiology, 99(3):225–234.

Forstmann BU, Dutilh G, Brown S, Neumann J, Von Cramon DY, Ridderinkhof KR, and Wagenmakers EJ (2008). Striatum and pre-SMA facilitate decision-making under time pressure. Proceedings of the National Academy of Sciences, 105(45):17538–17542.

Gelman A, Carlin JB, Stern HS, Dunson DB, Vehtari A, and Rubin DB (2013). Bayesian Data Analysis. CRC press.

Gelman A and Rubin DB (1992). Inference from iterative simulation using multiple sequences. Statistical Science, 7:457–472.

Gold JI and Shadlen MN (2007). The neural basis of decision making. Annual Review of Neuroscience, 30:535–574.

Heekeren HR, Marrett S, Bandettini PA, and Ungerleider LG (2004). A general mechanism for perceptual decision-making in the human brain. Nature, 431(7010):859–862.

Heitz RP (2014). The speed-accuracy tradeoff: history, physiology, methodology, and behavior. Frontiers in neuroscience, 8:150.

Heitz RP and Schall JD (2012). Neural mechanisms of speed-accuracy tradeoff. Neuron, 76(3):616–628.

Ivanoff J, Branning P, and Marois R (2008). fMRI evidence for a dual process account of the speed-accuracy tradeoff in decision-making. PLoS one, 3(7): e2635.

Karahan E, Costigan AG, Graham KS, Lawrence AD, and Zhang J (2019). Cognitive and white-matter compartment models reveal selective relations between corticospinal tract microstructure and simple reaction time. Journal of Neuroscience, 39(30):5910–5921.

Krummenacher J, Müller HJ, and Heller D (2002). Visual search for dimensionally redundant pop-out targets: Redundancy gains in compound tasks. Visual Cognition, 9(7):801–837.

Kruschke J (2014). Doing bayesian data analysis: A tutorial with r, jags, and stan.

Levitt H (1971). Transformed up-down methods in psychoacoustics. The Journal of the Acoustical Society of America, 49(2B):467—477.

Liu CC and Watanabe T (2012). Accounting for speed—accuracy tradeoff in perceptual learning. Vision research, 61:107—114.

Loughnane GM, Newman DP, Bellgrove MA, Lalor EC, Kelly SP, and O’Connell RG (2016). Target selection signals influence perceptual decisions by modulating the onset and rate of evidence accumulation. Current Biology, 26(4):496—502.

Luck SJ and Hillyard SA (1994). Spatial filtering during visual search: evidence from human electrophysiology. Journal of Experimental Psychology: Human Perception and Performance, 20(5):1000.

Mazurek ME, Roitman JD, Ditterich J, and Shadlen MN (2003). A Role for Neural Integrators in Perceptual Decision Making. Cerebral Cortex, 13(11):1257—1269.

Mulder MJ, Keuken MC, van Maanen L, Boekel W, Forstmann BU, and Wagenmakers EJ (2013). The speed and accuracy of perceptual decisions in a random-tone pitch task. Attention, Perception, & Psychophysics, 75(5):1048—1058.

Nunez MD, Gosai A, Vandekerckhove J, and Srinivasan R (2019). The latency of a visual evoked potential tracks the onset of decision making. Neuroimage, 197:93—108.

Palmer J (1995). Attention in visual search: Distinguishing four causes of a set-size effect. Current directions in psychological science, 4(4):118—123.

Palmer J, Huk AC, and Shadlen MN (2005). The effect of stimulus strength on the speed and accuracy of a perceptual decision. Journal of vision, 5(5):1—1.

Rajananda S, Lau H, and Odegaard B (2018). A random-dot kinematogram for web-based vision research. J. Open Res. Softw., 6.

Rangelov D and Mattingley JB (2020). Evidence accumulation during perceptual decisionmaking is sensitive to the dynamics of attentional selection. NeuroImage, 220:117093.

Ratcliff R (2002). A diffusion model account of response time and accuracy in a brightness discrimination task: Fitting real data and failing to fit fake but plausible data. Psychonomic Bulletin & Review, 9(2):278—291.

Ratcliff R (2006). Modeling response signal and response time data. Cognitive psychology, 53(3):195—237.

Ratcliff R and McKoon G (2008). The diffusion decision model: theory and data for two-choice decision tasks. Neural computation, 20(4):873—922.

Ratcliff R and Tuerlinckx F (2002). Estimating parameters of the diffusion model: Approaches to dealing with contaminant reaction times and parameter variability. Psychonomic bulletin & review, 9(3):438—481.

Reynolds JH and Chelazzi L (2004). Attentional modulation of visual processing. Annual Review of Neuroscience, 27(1):611–647. PMID: 15217345.

Roitman JD and Shadlen MN (2002). Response of neurons in the lateral intraparietal area during a combined visual discrimination reaction time task. Journal of neuroscience, 22(21):9475–9489.

Schmiedek F, Oberauer K, Wilhelm O, Süß HM, and Wittmann WW (2007). Individual differences in components of reaction time distributions and their relations to working memory and intelligence. Journal of Experimental Psychology: General, 136(3): 414.

Semmelmann K and Weigelt S (2017). Online psychophysics: Reaction time effects in cognitive experiments. Behavior Research Methods, 49(4):1241–1260.

Shadlen M, Britten K, Newsome W, and Movshon J (1996). A computational analysis of the relationship between neuronal and behavioral responses to visual motion. Journal of Neuroscience, 16(4):1486–1510.

Spiegelhalter DJ, Best NG, Carlin BP, and Van Der Linde A (2002). Bayesian measures of model complexity and fit. Journal of the royal statistical society: Series b (statistical methodology), 64(4):583–639.

Standage D, Blohm G, and Dorris MC (2014a). On the neural implementation of the speed-accuracy trade-off. Frontiers in Neuroscience, 8:236.

Standage D, Wang DH, and Blohm G (2014b). Neural dynamics implement a flexible decision bound with a fixed firing rate for choice: a model-based hypothesis. Frontiers in Neuroscience, 8:318.

Vandekerckhove J, Tuerlinckx F, and Lee MD (2011). Hierarchical diffusion models for two-choice response times. Psychological methods, 16(1):44.

Wagenmakers EJ, Love J, Marsman M, Jamil T, Ly A, Verhagen J, Selker R, Gronau QF, Dropmann D, Boutin B, et al. (2018). Bayesian inference for psychology. part ii: Example applications with jasp. Psychonomic bulletin & review, 25(1):58–76.

Wang XJ (2002). Probabilistic decision making by slow reverberation in cortical circuits. Neuron, 36(5):955–968.

Wendelken C, Ditterich J, Bunge SA, and Carter CS (2009). Stimulus and response conflict processing during perceptual decision making. Cognitive, Affective, & Behavioral Neuroscience, 9(4):434–447.

Wickelgren WA (1977). Speed-accuracy tradeoff and information processing dynamics. Acta psychologica, 41(1):67–85.

Wiecki T, Sofer I, and Frank M (2013). Hddm: Hierarchical bayesian estimation of the drift-diffusion model in python. Frontiers in Neuroinformatics, 7:14.

Wong KF and Wang XJ (2006). A recurrent network mechanism of time integration in perceptual decisions. Journal of Neuroscience, 26(4):1314–1328.

Wyart V, Myers NE, and Summerfield C (2015). Neural mechanisms of human perceptual choice under focused and divided attention. Journal of neuroscience, 35(8):3485–3498. 25716848[pmid].

Yamaguchi M, Crump MJ, and Logan GD (2013). Speed-accuracy trade-off in skilled typewriting: Decomposing the contributions of hierarchical control loops. Journal of Experimental Psychology: Human Perception and Performance, 39(3):678.

Zhang J and Bogacz R (2010). optimal decision making on the basis of evidence represented in spike trains. Neural Computation, 22:1113–1148.

Zhang J and Rowe JB (2014). Dissociable mechanisms of speed-accuracy tradeoff during visual perceptual learning are revealed by a hierarchical drift-diffusion model. Frontiers in Neuroscience, 8:69.

