## Supplementary material for "Imperfect integration: sensory congruency between multiple sources modulates selective decision-making processes": Supplemetal Material

Dominik Krzemiński, Jiaxiang Zhang

March 14, 2021

### Supplementary methods

#### The neural-mass model of perceptual decision

We used the two-state neural-mass model (Wong and Wang, 2006) in the following form:

$$\frac{dS_i}{dt} = -\frac{S_i}{\tau_S} + (1 - S_i)\gamma r(I_{\mathrm{syn},i}) \ , \tag{1}$$

$$r(I_{\mathrm{syn},i}) = \frac{aI_{\mathrm{syn},i} - b}{1 - \exp(-d(aI_{\mathrm{syn},i} - b))} \ , \tag{2}$$

$$\begin{cases} I_{\mathrm{syn,L}} = & J_{\mathrm{L,L}}S_{\mathrm{L}} + J_{\mathrm{L,R}}S_{\mathrm{R}} + I_{\mathrm{in,L}} + I_{\eta,\mathrm{L}} \\ I_{\mathrm{syn,R}} = & J_{\mathrm{R,R}}S_{\mathrm{R}} + J_{\mathrm{R,L}}S_{\mathrm{L}} + I_{\mathrm{in,R}} + I_{\eta,\mathrm{R}} \ , \end{cases} \tag{3}$$

$$\tau_\eta \frac{dI_{\eta,i}}{dt} = -I_{\eta,i} + \eta_t \sqrt{\tau_\eta}\sigma_\eta \ , \tag{4}$$

where the index $i = \mathrm{L}, \mathrm{R}$ refers to two neural populations selective for leftwards and rightwards choices. The state variable $S$ describes the NMDA gating variable (fraction of open gates). It can be shown that $S$ has a bijection mapping on pre-synaptic firing rates (Wong and Wang, 2006). $r$ describes the population firing rate function that depends on the synaptic input current $I_{\text{syn},i}$. The following values of the parameters were used: $a = 270(\text{VnC})^{-1}$, $b = 108$ Hz, $d = 0.154$ s, $\gamma = 0.641 \cdot 10^{-3}$ , $\tau_S = 100$ ms.

The synaptic input current $I_{\text{syn},i}$ combines recurrent inputs, mutual inputs, external inputs that relate to sensory information ($I_{\text{in},i}$, Equation 1 in the main text) and the noise current $I_{\eta,\text{i}}$. The following symmetric synaptic coupling parameters were used: $J_{\text{A,A}} = J_{\text{B,B}} = 0.2601$ nA and $J_{\text{A,B}} = J_{\text{B,A}} = 0.0497$ nA, $J_{\text{ext}} = 5.2 \cdot 10^{-4}$ nA·Hz$^{-1}$.

The noise current $I_{\eta,i}$ is integrated with $\tau_\eta = 2$ ms (time decay of AMPA receptor activation) and random variable sample from Normal distribution $\eta$. The variance of the noise factor is kept constant $\sigma_\eta = 0.01972$.

For simulations, we use standard Euler's integration method with a time step of 1 ms.

### Supplementary figures

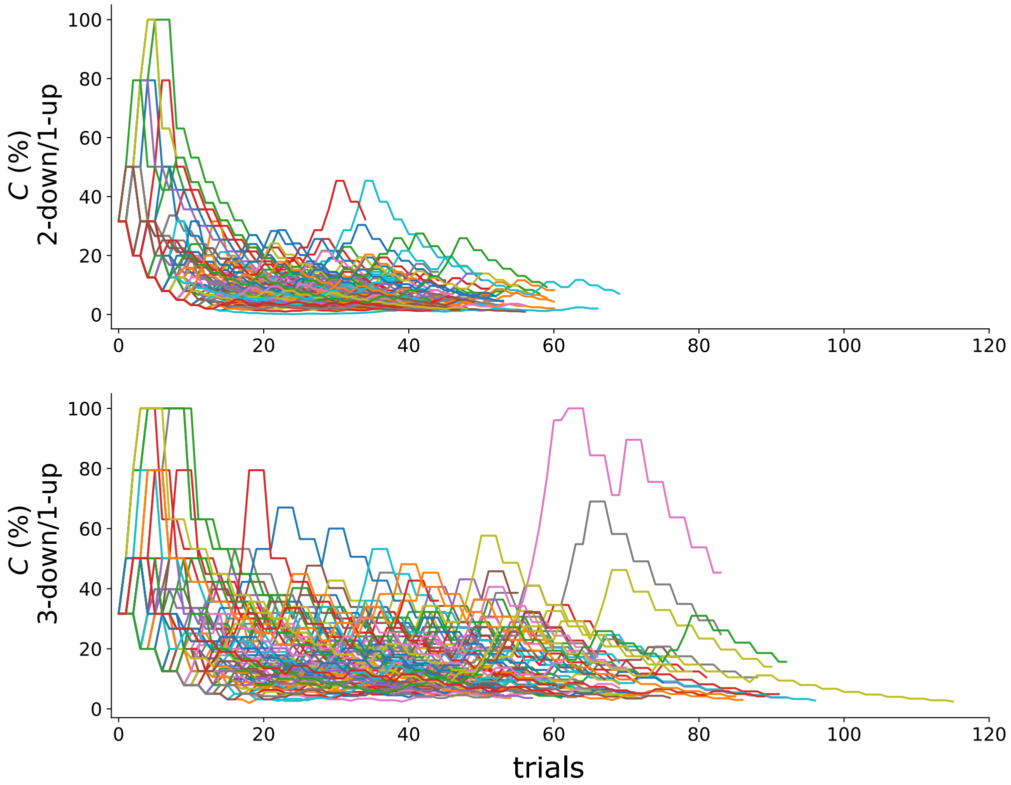


**Supplementary Figure 1:** Steps of the staircase procedure for the two staircase rules (top: two-up/one-down rule; bottom: three-up/one-down rule). Each line represents one participant.

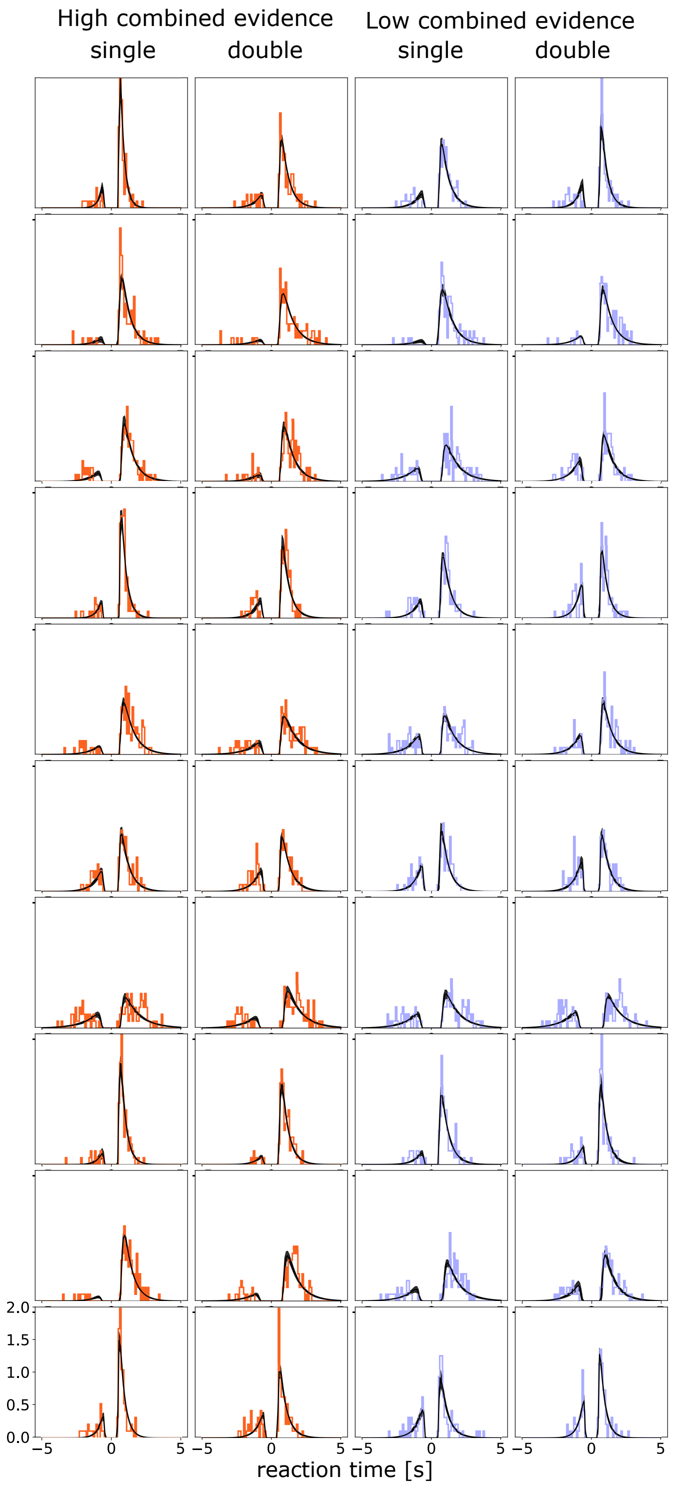


**Supplementary Figure 2:** Posterior predictive data distributions of 10 participants in Group 1 ($\theta = \pm 20°$). Each row shows data distributions (histograms) as well as Posterior model predictions (black lines) from the best fitted model from one of ten representative participants. The distributions along the positive x-axis indicate normalised correct response times, and the distributions along the negative x-axis indicate normalised error response times. Model predictions was generated in the same procedure as in Figure 6.

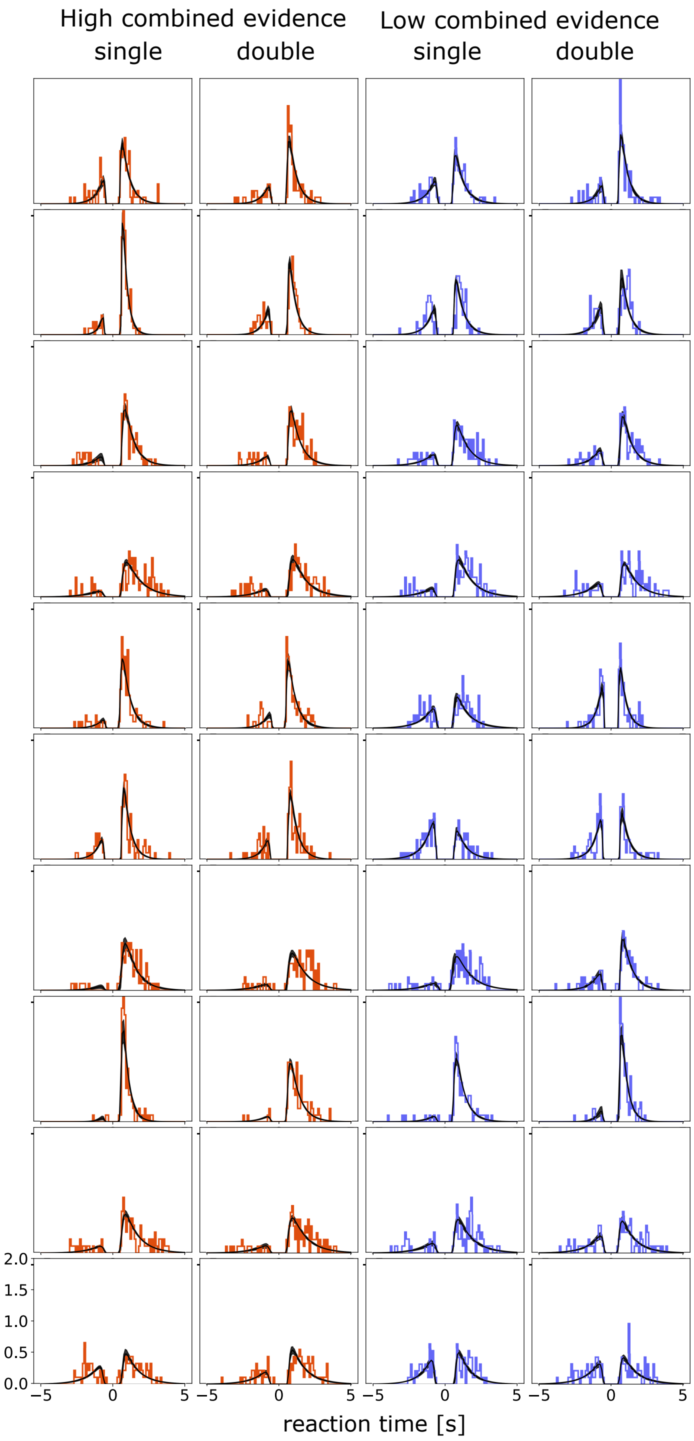


**Supplementary Figure 3:** Posterior predictive data distributions of 10 participants in Group 2 ($\theta = \pm 45°$). Each row shows data distributions (histograms) as well as Posterior model predictions (black lines) from the best fitted model from one of ten representative participants. The distributions along the positive x-axis indicate normalised correct response times, and the distributions along the negative x-axis indicate normalised error response times. Model predictions was generated in the same procedure as in Figure 6.

### References

Wong KF and Wang XJ (2006). A recurrent network mechanism of time integration in perceptual decisions. *Journal of Neuroscience*, 26(4):1314–1328.
